# Causal relations between cortical network oscillations and breathing frequency

**DOI:** 10.1101/2020.12.05.412999

**Authors:** Adriano BL Tort, Maximilian Hammer, Jiaojiao Zhang, Jurij Brankačk, Andreas Draguhn

**Author notes:** Correspondence (A.B.L. Tort) or (A. Draguhn). Equal contribution (Adriano BL Tort, Maximilian Hammer).

## Abstract

Nasal breathing generates a rhythmic signal which entrains cortical network oscillations in widespread brain regions on a cycle-to-cycle time scale. It is unknown, however, how respiration and neuronal network activity interact on a larger time scale: are breathing frequency and typical neuronal oscillation patterns correlated? Is there any directionality or causal relationship? To address these questions, we recorded field potentials from the posterior parietal cortex of mice together with respiration during REM sleep. In this state, the parietal cortex exhibits prominent theta and gamma oscillations while behavioral activity is minimal, reducing confounding signals. We found that the instantaneous breathing rate strongly correlates with the instantaneous frequency and amplitude of both theta and gamma oscillations. Granger causality analysis revealed specific directionalities for different rhythms: changes in theta activity precede and cause changes in breathing rate, suggesting control of breathing frequency by the functional state of the brain. On the other hand, the instantaneous breathing rate Granger-causes changes in gamma oscillations, suggesting that gamma is influenced by a peripheral reafference signal. These findings show that breathing causally relates to different patterns of rhythmic brain activity, revealing new and complex interactions between elementary physiological functions and neuronal information processing.

**Significance Statement:** The study of the interactions between respiration and brain activity has been focused on phase-entrainment relations, in which cortical networks oscillate phase-locked to breathing cycles. Here we discovered new and much broader interactions which link respiration rate (frequency) to different patterns of oscillatory brain activity. Specifically, we show that the instantaneous breathing rate strongly correlates with the instantaneous frequency and amplitude of theta and gamma oscillations, two major network patterns associated with cognitive functions. Interestingly, causality analyses reveal that changes in breathing rate follow theta, suggesting a central drive, while in contrast, gamma activity follows changes in breathing rate, suggesting the role of a reafferent signal. Our results reveal new mechanisms by which nasal breathing patterns may influence brain functions.

## Introduction

Several recent studies have shown that nasal breathing induces local field potential (LFP) oscillations at the same frequency as respiration in multiple regions of the rodent and human brain (for reviews, see Tort et al., 2018a; Heck et al., 2019). LFP phase-locking to respiratory cycles includes regions not primarily related to olfaction (Ito et al., 2014; Lockmann et al., 2016; Nguyen Chi et al., 2016; Zelano et al., 2016; Biskamp et al., 2017; Herrero et al., 2018; Karalis and Sirota, 2018; Kőszeghy et al., 2018; Rojas-Líbano et al., 2018; Tort et al., 2018b), suggesting that respiration-entrained oscillations aid the integration of widespread information (Heck et al., 2017, 2019; Tort et al., 2018a), similar to the proposed function of other slow network rhythms (Isomura et al., 2006; Canolty and Knight, 2010). Given the alleged beneficial effects of some respiratory practices to mood, behavioral performance, and cognitive abilities (Pascoe and Bauer, 2015; Zelano et al., 2016; Melnychuk et al., 2018; Nakamura et al., 2018; Perl et al., 2019; Novaes et al., 2020), the fact that nasal respiration can modulate brain oscillations in non-olfactory regions has sparked large interest, including in popular science media, since it could provide a way through which respiration would affect brain functions (Heck et al., 2017; Varga and Heck, 2017; Tort et al., 2018a).

The research on relations between respiration and neuronal network activity has so far mainly focused on phase-entrainment relations, in which a given cortical LFP is shown to oscillate phase-locked to breathing cycles, typically accompanied by a peak in the LFP power spectrum at the breathing rate (Tort et al., 2018b), or by modulation of action potential probability by the respiration phase (Zhong et al., 2017; Karalis and Sirota, 2018; Kőszeghy et al., 2018). Other studies have shown that the respiration phase is also capable of modulating the instantaneous amplitude of faster, gamma-frequency oscillations (Ito et al., 2014; Yanovsky et al., 2014; Rojas-Líbano et al., 2018; Cavelli et al., 2019), especially in frontal regions (Biskamp et al., 2017; Zhong et al., 2017). All these findings are, however, confined to the time domain of individual oscillation cycles. Nevertheless, both breathing rate and the pattern of neuronal network oscillations vary over time, and it remains unclear whether there is any interaction between these rhythmic phenomena at the time scale of such variations. Of note, we have recently reported such an interaction by showing that the strength of theta-gamma coupling depends on respiratory frequency (Hammer et al., 2020).

We therefore performed a systematic analysis of relations between the varying patterns of cortical oscillations and breathing. In particular, we searched for correlations and causality between the instantaneous breathing rate and the instantaneous amplitude and frequency of neocortical network oscillations. To that end, we recorded respiration through a whole-body plethysmograph while simultaneously recording field potentials from the posterior parietal cortex of mice. We focused on periods of REM sleep, which is characterized by prominent activity in the theta and gamma bands (Montgomery et al., 2008; Scheffzük et al., 2011). Moreover, this state guarantees a stable behavior; that is, any observed changes in breathing are not due to changes in motor demands, and cognitive-behavioral interference with the environment is minimal. Our results reveal strong correlations between instantaneous breathing rate and instantaneous amplitude and frequency of theta and gamma oscillations. We also report a clear directionality, with theta oscillations being Granger-causal for changes in breathing frequency which, in turn, is causal for gamma activity.

## Results

### Respiration phase causes neocortical oscillations at the same frequency as breathing

We recorded parietal cortex activity of mice during REM sleep while mice were placed in a plethysmograph to simultaneously monitor their respiratory activity (Figure 1A). REM episodes were identified by behavioral immobility and prominent theta activity in the cortical field potential (for details, see Materials and Methods). The power spectrum of the plethysmograph-derived respiration signal exhibited a peak at 3.2 ± 0.35 Hz (mean ± SD, n=22 mice) with a long tail towards faster frequencies up to values ∼11 Hz. At the same time, the neocortical local field potential (LFP) displayed – as expected – a prominent peak at theta frequency (7.7 ± 0.37 Hz; Figure 1B). In addition, a smaller power bump at a similar frequency as the respiration power peak could also be observed in the LFP spectrum (Figure 1B); moreover, the phase coherence spectrum between respiration and LFP peaked at respiration frequency (Figure 1C). Therefore, combined, the power and coherence spectra show that – in addition to theta oscillations – the parietal cortex LFP also displays the so-called respiration-entrained rhythm (RR) during REM sleep, consistent with previous reports (Zhong et al., 2017; Tort et al., 2018b). Also consistent with these reports, there was no phase coherence between the parietal field potential and respiration at theta frequency (Figure 1C).

**Figure 1.**
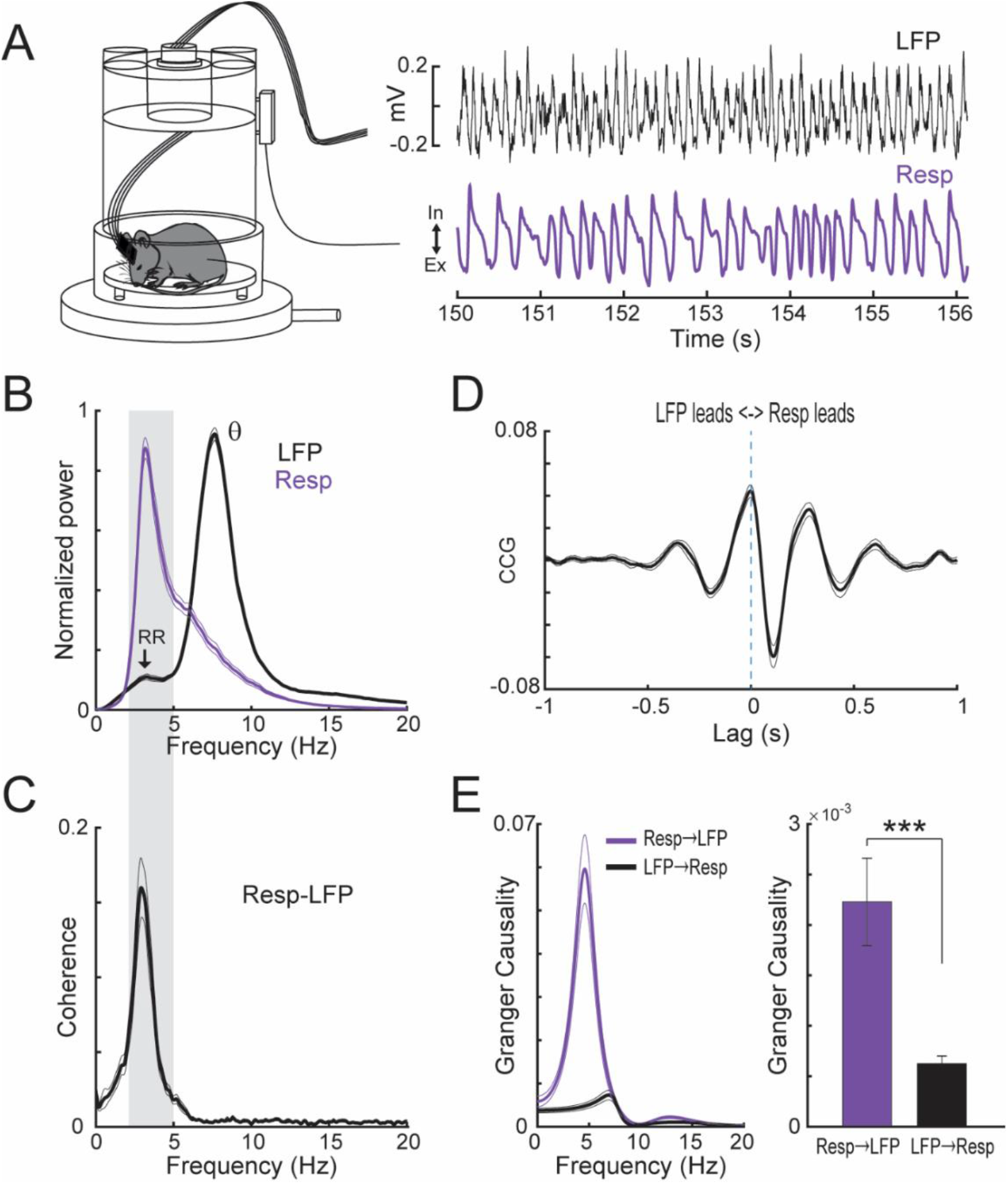
Respiration drives phase-entrained oscillations in the parietal cortex during REM sleep. (A) Depiction of a plethysmograph recording of respiration (Resp) along with the parietal LFP during REM sleep. (B) Mean (± SEM) power spectra for the two signals (n=22 mice; the individual spectra were normalized by the maximal value before computing the mean). In addition to prominent theta (θ, 6-12 Hz) oscillations, the LFP spectrum has a smaller power bump at a similar frequency range as respiration (2-5 Hz), which corresponds to the respiration-entrained LFP rhythm (RR; see Tort et al., 2018b). (C) Mean phase coherence spectrum between the respiration and LFP signals, showing marked coherence at the respiration (and not theta) frequency. (D) Mean cross-correlogram (CCG) between Resp and LFP. (E) Mean Granger causality for the Resp->LFP (blue) and LFP->Resp (black) directions. The left panel shows the frequency spectrum; the right panel shows the overall (time domain) causality levels of each direction. Respiration highly Granger-causes LFP activity at the breathing frequency (that is, respiration causes RR). Notice that the raw LFP and Resp signals were used in this analysis (as opposed to results shown in the subsequent figures). ***p<0.001.

After corroborating RR presence in the parietal cortex, we next characterized directionality relations between respiration and LFP signals. To that end, we computed both cross-correlograms (CCGs) and Granger causality between their raw traces (see Materials and Methods). Consistent with the phase coherence spectrum (Figure 1C), the CCGs exhibited an oscillatory pattern with a period matching the respiration cycle (321 ± 30 ms [mean ± SD; n=22 mice]; Figure 1D). Interestingly, using the respiration signal as reference, the average CCG exhibited a negative peak at 110 ms, indicating that the LFP phase entrainment to respiratory cycles lags the respiration signal (Figure 1D). It should be noted, though, that due to the rhythmical nature of the CCG in this case, directionality relations cannot be unambiguously determined (Bastos and Schoffelen, 2015). We therefore calculated the Granger causality spectrum between raw LFP and respiration signals, which revealed a prominent causality peak at breathing frequency specifically for the resp → LFP direction (Figure 1E left). Consistently, the overall, time-domain Granger causality was statistically significantly higher in this direction than in the LFP → resp direction (t(21)=4.06, p=0.00056, paired t-test; Figure 1E right). Together, these results corroborate previous observations that respiration drives neuronal network oscillations of the same frequency (Nguyen Chi et al., 2016; Karalis and Sirota, 2018).

### Theta frequency and amplitude depend on breathing rate

Respiration frequency is dynamically modulated (Figure 1A,B), similar to other, brain-endogenous rhythms such as theta or gamma oscillations. This poses the question of whether there is any influence of breathing rate on oscillatory network activity or vice versa. We therefore tracked the breathing rate during REM sleep by computing time-frequency spectrograms of respiratory activity using 500-ms windows with no overlap. For each window, to aid visualization the power distribution was normalized by its maximal value, and the instantaneous breathing rate was defined as the frequency of the maximal power value (which equals 1 due to the normalization; Figure 2A,B). The breathing rate typically varied between 2 and 5 Hz during the analyzed REM sleep periods, though faster breathing up to 11 Hz could also be observed (Figure 2A-C). A similar procedure was performed to track the peak frequency of theta in the parietal cortex LFP signal for the same 500-ms windows. Interestingly, the histogram distribution of respiration and theta peak frequencies for the pool of 500-ms windows (Figure 2D) resembled the average power spectra (c.f. Figure 1B). This shows that the respiration power tail towards higher frequencies (Figure 1B) corresponds to periods in which the animals indeed breathed at faster rates, as opposed to being caused by power leakage from lower breathing frequencies.

**Figure 2.**
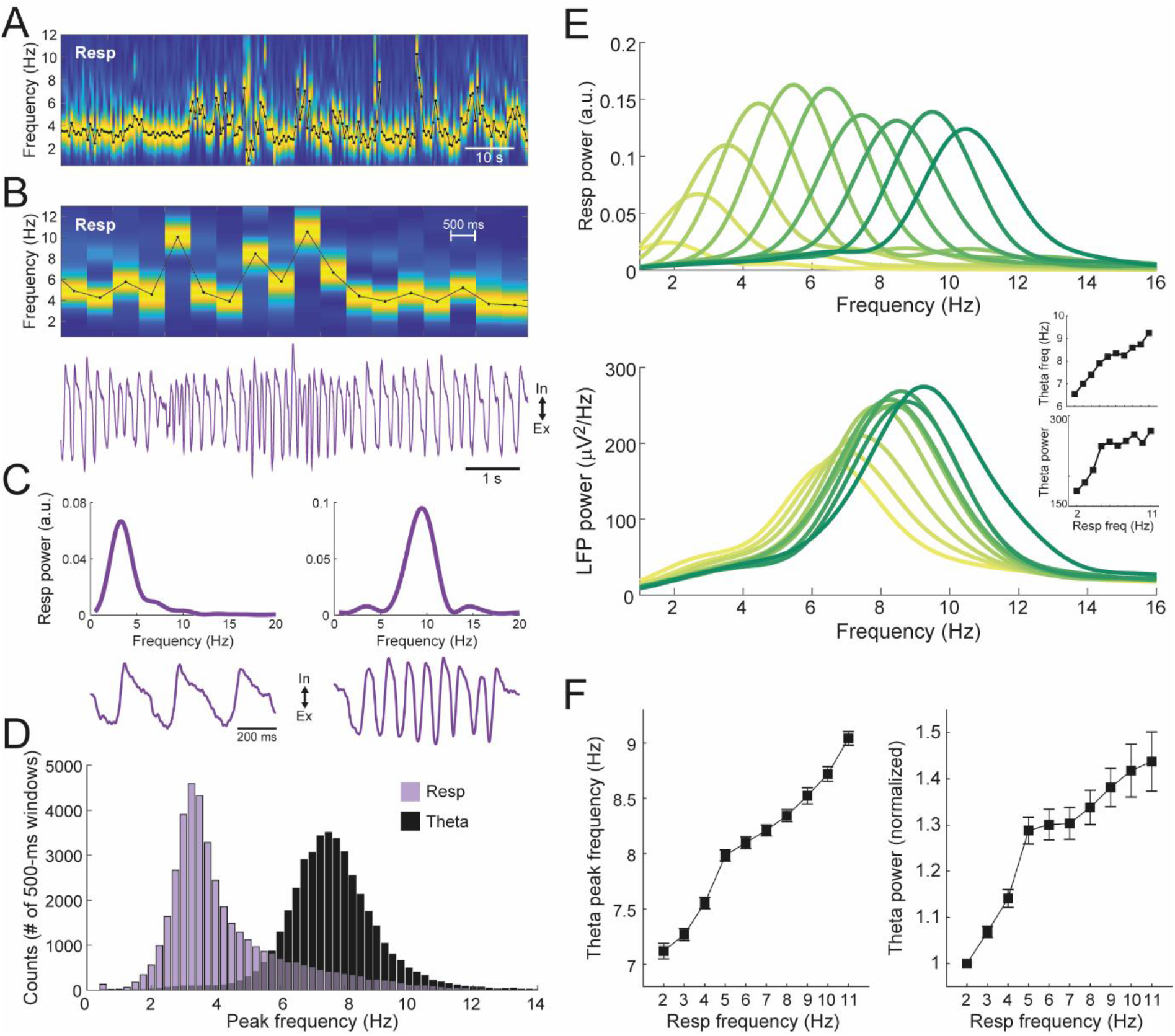
Tracking respiration during REM sleep reveals a link between breathing rate and theta activity. (A) Example time-frequency power spectrogram for the Resp signal using non-overlapping 500-ms windows. The power spectrum of each window was normalized by the maximal value. The black line depicts the breathing rate, as defined by the peak power frequency. (B) Shown are 9 seconds of a Resp spectrogram (top) and the corresponding raw signal (bottom). (C) Example of Resp power spectra (top) and raw signals for two 1-s epochs differing in breathing rate. (D) Histogram counts of theta and respiration peak frequency in 500-ms windows (pool over 22 animals). (E) (Top) Mean Resp power spectra for (overlapping) subsets of 500-ms windows binned by breathing rate (respiration frequency intervals: 1-3 Hz, 2-4 Hz, 3-5 Hz, 4-6 Hz, 5-7 Hz, 6-8 Hz, 7-9 Hz, 8-10 Hz, 9-11 Hz, and >10 Hz). Results obtained for a representative animal. (Bottom) Average LFP power spectra for the same respiration-binned time windows, as labeled by the color code. Notice that both theta power and theta frequency increase with faster breathing. The inset plots show theta peak frequency and power as a function of respiration frequency. (F) Group data of theta frequency and normalized theta power as a function of respiration frequency (mean ± SEM; n=22 mice). The normalization was performed within animals by dividing power values by the value in the first respiration frequency bin.

We next grouped time windows with similar instantaneous breathing frequency (Figure 2A,B) into ten groups (2-Hz bandwidth with 1-Hz overlap, that is: 1-3 Hz, 2-4 Hz, 3-5 Hz, etc). Figure 2E shows power spectra from data for each instantaneous frequency (top) together with the mean LFP power spectra for the same “respiration-binned” window subsets (bottom) in one animal. This alignment revealed a major increase in both theta frequency and theta power with increasing breathing rate (Figure 2E bottom). This result was very robust at the group level of 22 mice, for which repeated measures ANOVA showed a highly significant effect of respiration rate on both theta frequency and power (Figure 2F; theta frequency: F(9,203)=107.32, p<10^−71^; theta power: F(9,203)=17.27, p<10^−20^). Therefore, in addition to the previously described cycle-by-cycle phase-locking effects (Figure 1), breathing does also relate to oscillatory brain activity through its instantaneous rate (Figure 2F).

### Changes in theta activity precede changes in breathing rate

The results in Figure 2F show that both, instantaneous frequency and amplitude of theta exhibit a strong positive relationship with the instantaneous breathing rate. In order to assess potential causal relations, we next investigated directionality between these oscillatory features. To that end, we performed a similar approach as above, but this time, to allow for CCG and Granger causality calculations, we used a 90% overlap among the 500-ms windows (i.e., 50-ms step size). Therefore, for the directionality analyses we used time series of instantaneous frequency and/or power sampled at 20 Hz. Surprisingly, the average CCG computed using the breathing rate as the time lag reference showed that the changes in theta frequency occurred before the changes in breathing rate (−125 ± 53 ms [mean ± SD], t(21)=−11.08, p<10^−9^, one sample t-test against 0 ms; Figure 3A). This finding was corroborated by Granger causality analysis in the frequency and time-domain (Figure 3B), in which causality in the direction theta freq → resp freq was statistically significantly higher than in the opposite direction (t(21)=−4.77, p=0.0001, paired t-test). Thus, the instantaneous frequency of theta oscillations is Granger-causal for the instantaneous frequency of respiration. We note that this result appears to contrast the opposite Granger-causality between the raw respiration and parietal cortex LFP signals shown in Figure 1E. Here, however, we analyzed directionality relations between the time series of instantaneous frequencies, a derived feature from the raw signals reflecting the variance in leading frequency, rather than the actual oscillation itself. In line with this notion, the high Granger causality at slow frequencies in the left panel of Figure 3B does not correspond to respiration- or theta-frequencies but is likely due to the slower time-scale on which variations of theta and breathing occur.

**Figure 3.**
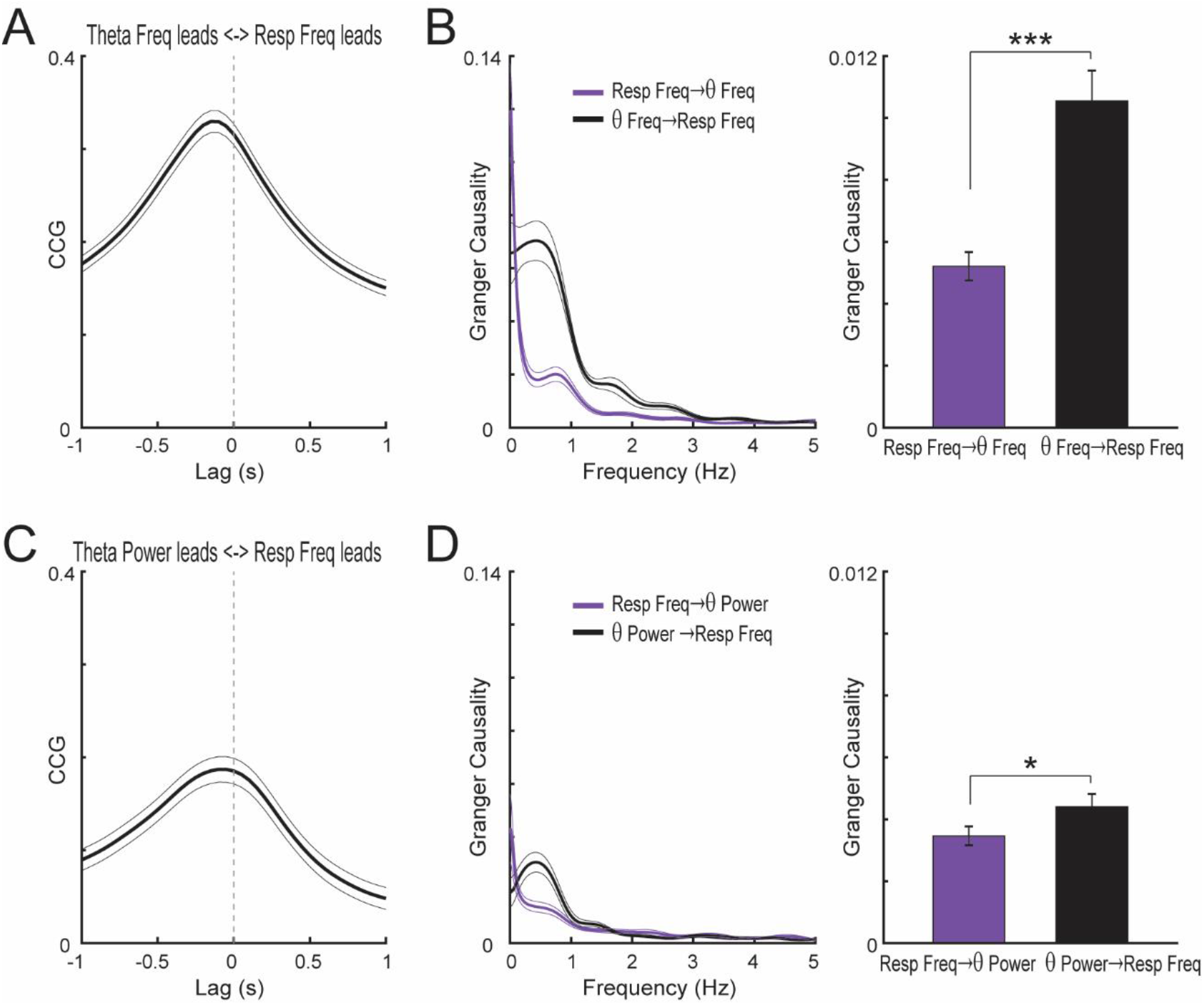
Changes in theta frequency precede changes in breathing rate. (A) Mean CCG (± SEM) between the time series of the instantaneous frequency of theta and respiration during REM sleep (n=22 mice). (B) Frequency- (right) and time-domain (left) Granger-causality between of the instantaneous frequency of theta and respiration, showing that theta frequency changes Granger-causes breathing rate changes during sleep. (C,D) As above, but for theta power. For these and subsequent directionality analyses, the peak LFP frequency and power and the peak Resp frequency were estimated using overlapping 500-ms windows (see Methods for details). *p<0.05; ***p<0.001.

We then asked whether variations in theta power would also be causally related to respiration. Using the time series of instantaneous breathing rate as reference, we found that theta power changes tended to lead respiration rate, as inferred by both the mean CCG lag (−80 ± 10 ms [mean ± SD], t(21)=−3.54, p=0.0019, one sample t-test against 0 ms; Figure 3C) and Granger causality (t(21)=−2.12, p=0.046, paired t-test between time domain Granger causality levels in each direction; Figure 3D). The magnitude of this effect was, however, much weaker than the directionality of variations in frequency (t(21)=8.11, p<10^−7^, paired t-test between Granger causality levels). In all, as opposed to the phase-entrainment relations which are driven by respiration (Figure 1), these results show that the instantaneous breathing rate does not lead but actually follows intrinsic changes in brain activity in the theta range.

### Gamma frequency increases with breathing rate

In addition to RR and theta oscillations (Figure 1B), the parietal cortex was also characterized by activity in two gamma sub-bands during REM sleep. These could be clearly inferred by the power bumps at ∼50 to ∼80 Hz and ∼120 to ∼160 Hz in the LFP spectrum, especially when plotted at a logarithmic scale (Figure 4A left). In this work, we refer to these frequency ranges as slow- and fast-gamma activity, respectively.

**Figure 4.**
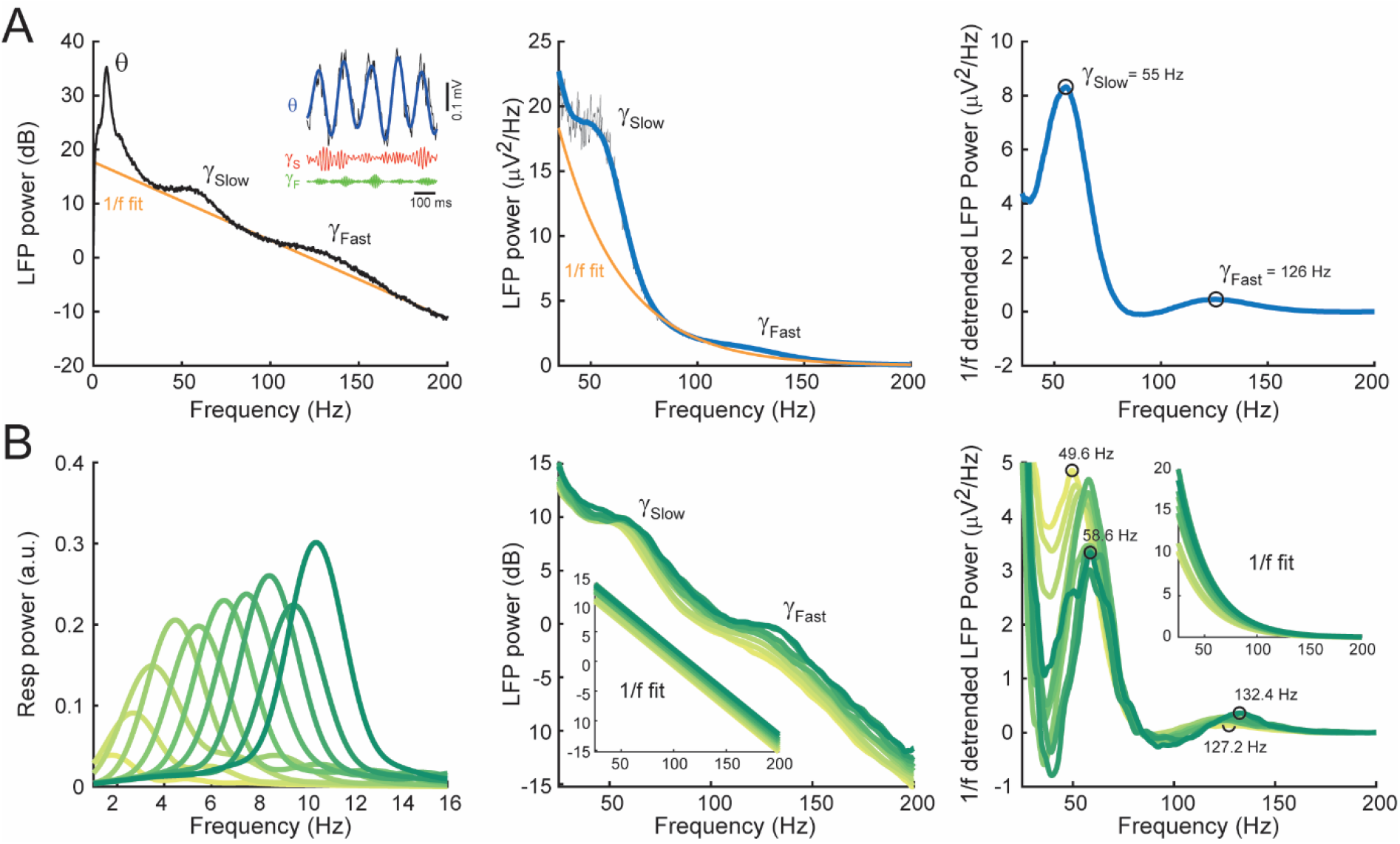
Estimating gamma frequency and power. (A) (Left) Average LFP power spectrum in logarithm (dB) scale. Notice power bumps at the slow and fast gamma sub-bands. The 1/f power fit is shown in orange (see Materials and Methods). Inset shows example raw (black) and filtered (colored, as labeled) LFP traces. (Middle) Power values of fast LFP frequencies (40-200 Hz) in linear scale (gray trace). The thick blue curve shows smoothed power values. (Right) 1/f-detrended power spectrum, obtained by subtracting the 1/f fit from the smoothed power. The peak frequency of slow and fast gamma was estimated from the detrended power curve. (B) (Left) Mean Resp power spectra for respiration-binned time windows as in Figure 2E (different animal). (Middle) Mean LFP power spectra for the same respiration-binned time windows plotted in logarithmic scale and focused in the fast frequency bands. The inset plot shows the 1/f fits. The frequency and absolute power of both gamma sub-bands increase with faster breathing. (Right) Mean 1/f-detrended power spectra. Notice a clear decrease in detrended slow gamma power with increasing breathing rate.

We first call attention to a technical point that will influence the interpretation of the results: in order to properly track the activity of genuine gamma oscillations – as opposed to unspecific activity in the gamma range – we estimated gamma frequency and power for each sub-band after first subtracting the 1/f fit from the LFP power spectrum (see Figure 4A,B, right panels). Therefore, by “genuine” activity we mean the excess power in the analyzed gamma sub-band above the 1/f power decay level. This approach controls for broad, unspecific changes of overall power at fast frequencies which can occur, for example, due to EMG contamination or other sources of noise. In any case, to allow comparison with previous approaches (e.g., Chen et al., 2011; Zheng et al., 2015), in Figure S1 we also present results obtained from un-subtracted power spectra, irrespective of the presence or not of prominent gamma power bumps.

Figure 4B shows power spectra computed for REM sleep epochs binned by respiration frequency as in Figure 2E, but for another example animal. The middle panel shows the absolute LFP power and the right panel shows the 1/f-detrended power. Notice in these panels an increase in the peak frequency of both slow- and fast-gamma oscillations for epochs associated with faster breathing rates (dark green). The relation between slow- and fast-gamma power with breathing rate, on the other hand, depended on how gamma power was estimated. Considering the absolute power level (Figure 4B middle), there was a clear increase during faster breathing for both gamma sub-bands. However, this apparent power increase did not seem to be specific for the gamma sub-bands since there was a broadband shift towards higher power levels for all fast frequencies. In fact, this was reflected in the upshift also observed for the 1/f power fits (see insets in Figure 4B). When removing this general trend by subtracting the corresponding 1/f fit of each spectrum, it became evident that the genuine slow-gamma power decreased at faster breathing (Figure 4B right), while the picture for genuine fast-gamma power was less clear.

A systematic analysis of group data (Figure 5) revealed that the increase of gamma frequency with breathing rate was highly statistically significant for both sub-bands (slow-gamma frequency: F(9,203)=4.23, p<0.0001; fast-gamma frequency: F(9,203)=3.42, p=0.0006, repeated measures ANOVA), and so was the decrease in genuine slow-gamma power (Figure 5A right; F(9,203)=6.96, p<10^−8^). On the other hand, genuine fast gamma power exhibited large variability among animals and was not statistically significantly related to breathing rate (Figure 5B right; F(9,203)=59.31, p=0.88). In summary, these data show an interdependence between slow and fast gamma oscillations and respiration rate, independent from a general increase in high-frequency LFP-components at higher breathing frequencies (see Figures S1 to S3).

**Figure 5.**
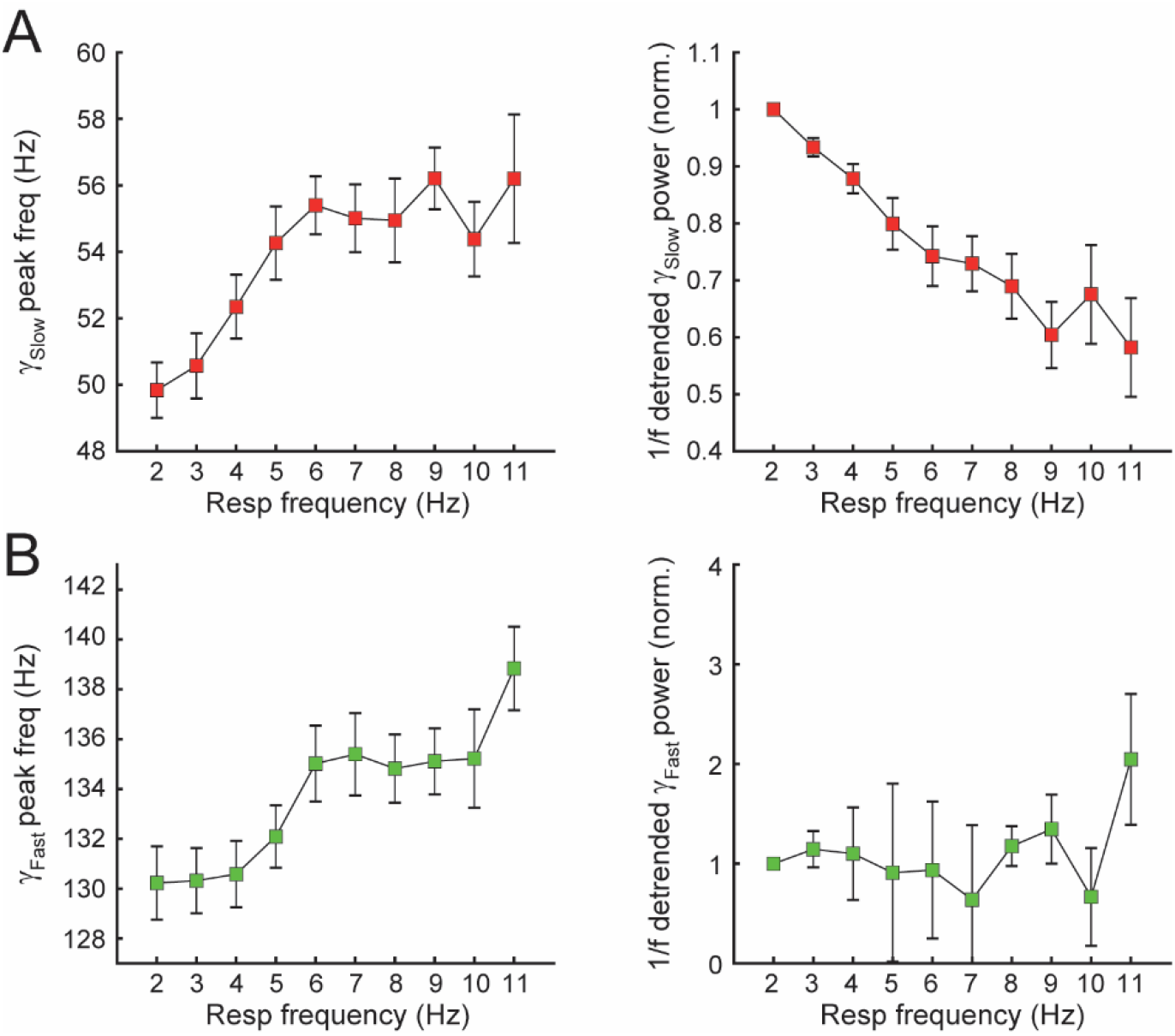
Gamma activity depends on breathing rate. (A,B) Group data of peak frequency (left) and 1/f-detrended power (right) as a function of respiration frequency (mean ± SEM; n=22 mice) for slow (A) and fast (B) gamma. Power values were normalized by the first respiration frequency bin.

### Changes in slow-gamma frequency and power follow changes in breathing rate

Similar to theta oscillations, we next analyzed directionality relations between breathing rate and the instantaneous frequency of both gamma sub-bands. For slow-gamma oscillations, using respiration as reference, the CCG peaked at 168 ± 13 ms (mean ± SD), meaning that the increases in instantaneous slow-gamma frequency lagged the increases in breathing rate (t(21)=6.13, p<10^−5^, one sample t-test against 0 ms; Figure 6A top). Granger causality analysis revealed a consistent finding, in which causality levels in the respiration → slow-gamma direction were statistically significantly higher than in the slow-gamma → respiration direction (Figure 6B top; t(21)=10.25, p<10^−8^, paired t-test). For the fast-gamma band, the CCG exhibited a peak at −109 ± 174 ms (t(21)=−2.95, p=0.008, one sample t-test against 0 ms; Figure 6A bottom), suggesting a possible lead by fast-gamma, though the Granger analysis showed a tendency for higher causality in the respiration → fast-gamma than in the fast-gamma → respiration directions (Figure 6B bottom; t(21)=1.78, p=0.09, paired t-test). In any case, Granger causality levels for respiration and fast-gamma frequency were much lower than the causality levels found between respiration and slow-gamma frequency (t(21)=−8.07, p<10^−7^, paired t-test).

**Figure 6.**
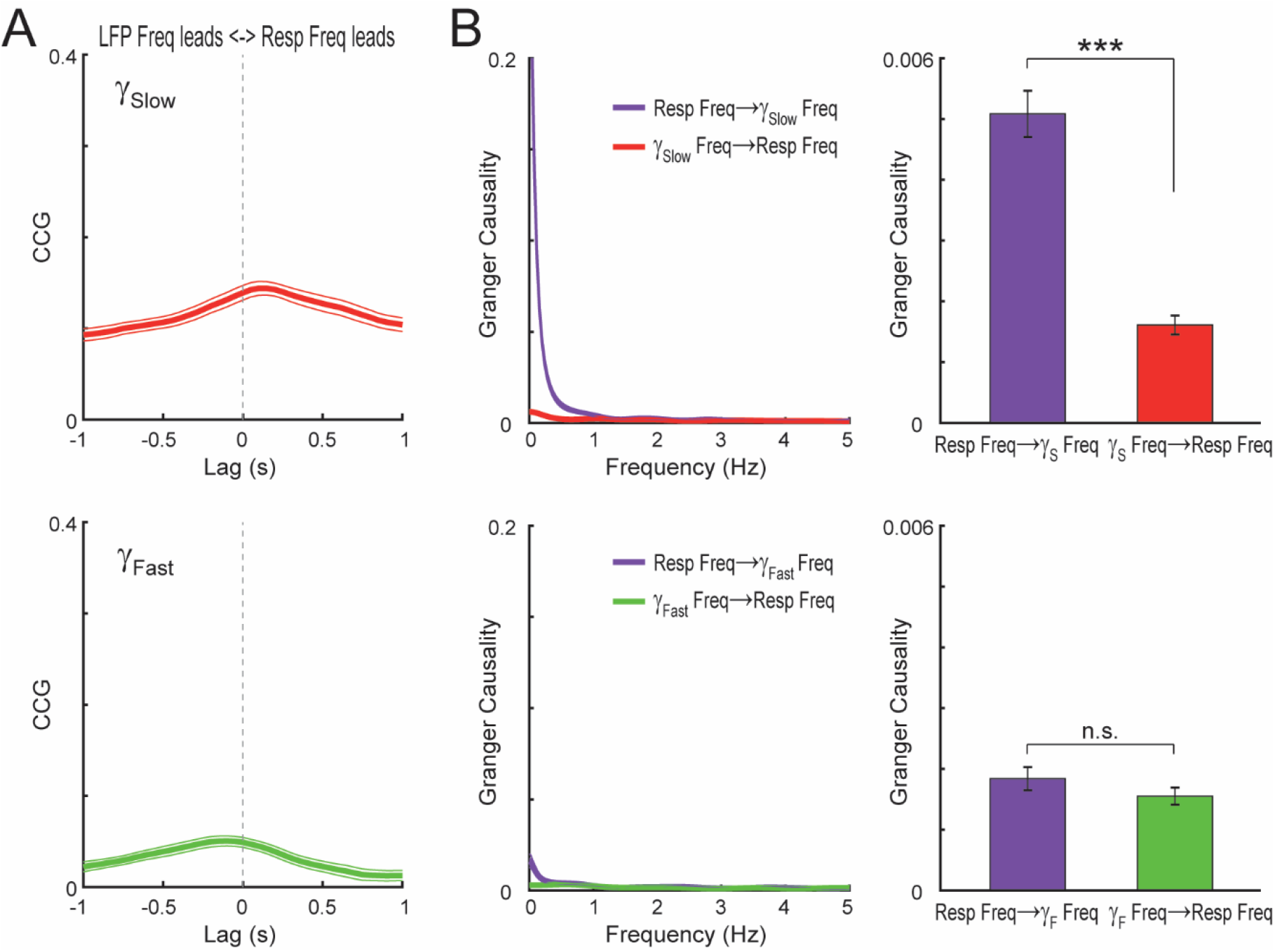
Changes in slow gamma frequency follow changes in breathing rate. (A) Mean CCG (± SEM) between the instantaneous frequency of slow (top) and fast (bottom) gamma frequencies and respiration rate during REM sleep (n=22 mice). (B) Frequency- (right) and time-domain (left) Granger-causality between the instantaneous frequency of gamma and respiration. Respiration frequency Granger-causes slow gamma frequency while having no directionality relation with fast gamma frequency. ***p<0.001.

Finally, we investigated directionality relations between gamma power and breathing rate. For the absolute power values (Figure S1), we found that the breathing rate was Granger-caused by both slow-gamma and fast-gamma power (slow gamma: t(21)=−6.07, p<10^−5^; fast gamma: t(21)=−9.82, p<10^−8^, paired t-tests between directions), a result confirmed by inspection of the CCG peak lags (slow gamma: −175 ± 72 ms [mean ± SD], t(21)=−11.41, p<10^−9^; fast gamma: −120 ± 45 ms, t(21)=−12.44, p<10^−10^, one sample t-tests against 0 ms; Figure S1). However, we did not take this result at face value because some corollaries called our attention. For one, it sounded odd that slow-gamma power would drive the breathing rate while its frequency would follow (c.f. Figure 6). For another, it also seemed strange that causality levels were much higher for the fast-gamma band, which usually has the lowest signal-to-noise ratio; in fact, fast-gamma causality was even higher than theta power causality levels (c.f. Figure 3D). We thus suspected that these causality relations could be due to unspecific variations in fast frequency activity. Consistent with this possibility, we found even higher causality levels for the very fast frequency range of 180-230 Hz (Figures S2 and S3), which had no power peak whatsoever and – to the best of our knowledge – is non-physiological during REM sleep. Interestingly, and consistent with a possible role of LFP contamination by extra brain sources, we found that the accelerometer signals were even more causal to respiration rate than fast LFP power (Figures S2 and S3). Thus, we conclude that the apparent causality between uncorrected gamma power and respiration frequency is likely caused by increased muscular activity during faster breathing which contaminates the fast frequency components of the LFP spectrum.

The picture was much different when we analyzed the 1/f-corrected power levels (Figure 7). In this case, the directionality relations were qualitatively similar to those found for the instantaneous frequency: namely, genuine slow gamma power changes followed changes in breathing rate (CCG lag: 84 ± 155 ms [mean ± SD], t(21)=2.55, p=0.019, one sample t-test against 0 ms; Granger causality: t(21)=3.94, p=0.0007, paired t-test between directions; Figure 7A,B top), while fast gamma power was not causally related to breathing rate (CCG lag: 48 ± 290 ms, t(21)=0.77, p=0.45, one sample t-test against 0 ms; Granger causality: t(21)=−0.34, p=0.74, paired t-test between directions; Figure 7A,B bottom). Interestingly, notice that resp-slow gamma CCG is negative (Figure 7A top), consistent with the inverse relationship between genuine slow gamma power and respiration found in Figure 5A (right panel). In all, changes in respiration frequency precede and Granger-cause changes in slow- and fast-gamma frequency as well as slow-gamma power after correcting for broad, frequency-unspecific power shifts. This finding contrasts with the opposite temporal and causal relation found between respiration and theta, in which the LFP theta activity leads respiration (c.f. Figure 3).

**Figure 7.**
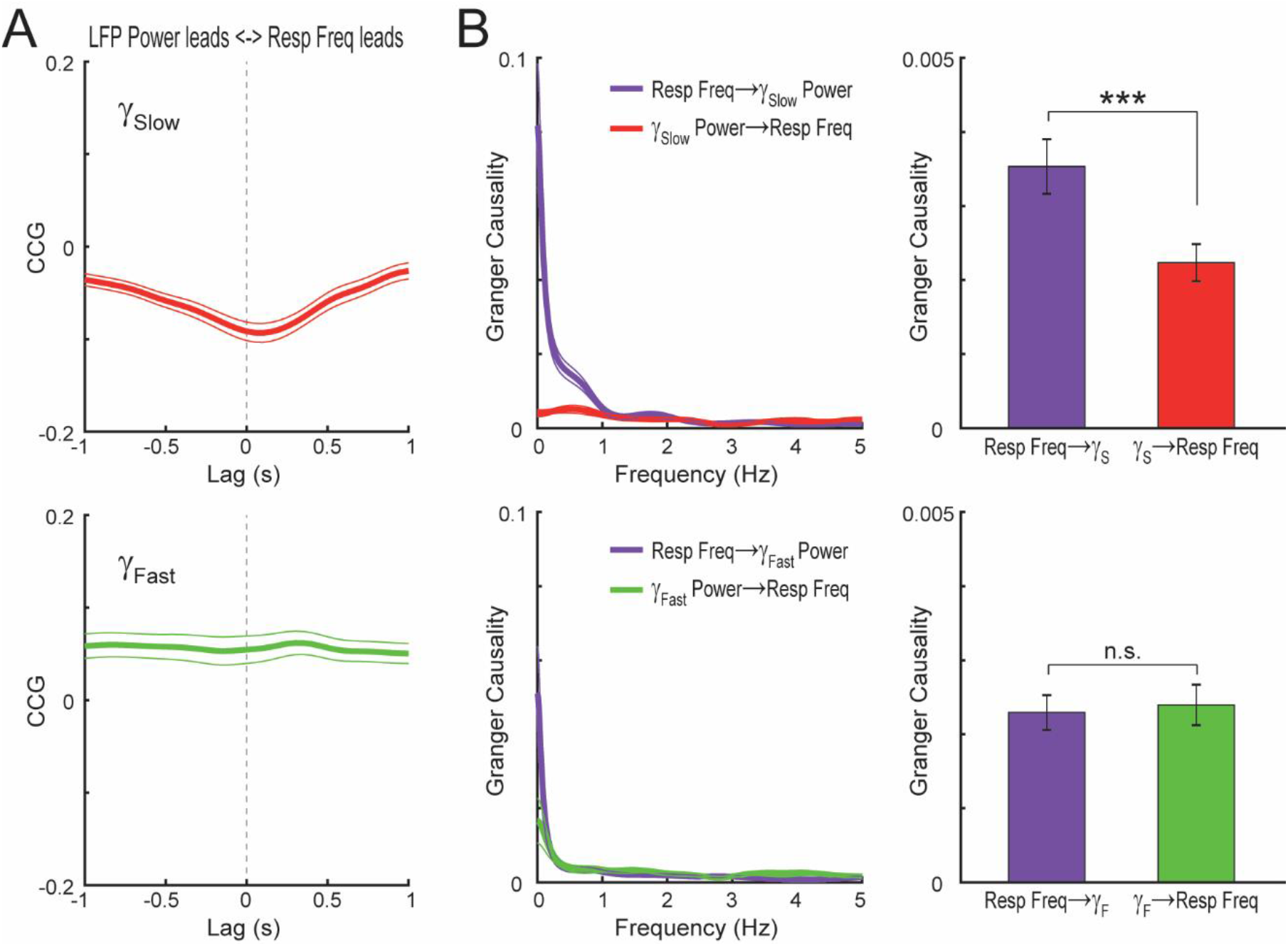
Slow gamma power (1/f-corrected) follows respiration rate. (A) Mean CCG (± SEM) between the 1/f-detrended power of slow (top) and fast (bottom) gamma and respiration rate during REM sleep (n=22 mice). (B) Frequency- (left) and time-domain (right) Granger-causality between 1/f-corrected gamma power and respiration. Respiration frequency Granger-causes slow gamma power and has no directionality relation with fast gamma power, thus similar to its effect over gamma frequency (c.f. Figure 6). ***p<0.001.

Combined, therefore, our result suggests that changes in the frequency of cortical theta oscillations precede and cause changes in respiration frequency (Figure 3). In contrast, the real-time respiration-entrained LFP component (Figure 1D,E) and slow-gamma activity (Figures 6 and 7) depend on, and follow nasal breathing.

## Discussion

The effect of respiration on brain activity has been the subject of much recent work. It is by now clear that respiration adds a breathing-frequency component to neuronal network oscillations (for reviews, see Tort et al., 2018a; Heck et al., 2019). Importantly, this effect occurs in widespread brain regions, far beyond olfactory networks, a finding reproduced here (Figure 1). The present work reveals a new, much broader interaction between respiration and different network oscillations. Both activities do not only interact through cycle-by-cycle phase-entrainments but also through mutual dependencies involving the instantaneous frequency. Specifically, we show that the frequency and amplitude of theta and gamma oscillations detected in the neocortex co-vary with breathing rate. Moreover, our directionality analyses revealed coherent temporal relations among the electrophysiological and respiratory rhythms, in which changes in theta activity precede changes in breathing rate that, in turn, precede changes in slow gamma. Therefore, the present study unveils novel relationships between respiration and brain activity beyond the well-established phase-locking relations.

A myriad of recent studies convincingly showed that the reafferent sensory signal brought about by nasal breathing drives LFP oscillations phase-locked to respiration (RR). This has been demonstrated in multiple ways, including experimental (e.g., naris closure, olfactory epithelium ablation, bulbectomy and tracheotomy; Ito et al., 2014; Yanovsky et al., 2014; Lockmann et al., 2016; Bagur et al., 2018; Karalis and Sirota, 2018; Moberly et al., 2018) and computational approaches (e.g., Granger causality analysis; Nguyen Chi et al., 2016; Karalis and Sirota, 2018). Here we show different relations between variations in instantaneous frequency of network oscillations and respiration. Indeed, cortical LFP activity in the theta range Granger-causes respiration frequency, rather than vice versa. Nevertheless, we believe that this result is consistent with a causal role of a central drive underlying changes in both network and breathing activity. Our working hypothesis, illustrated in Figure 8, is that REM-controlling neuronal circuits in the brainstem send signals to theta-generator circuits and to nearby respiration-controlling nuclei. Thereby, they determine changes in theta-as well as respiration frequency during REM sleep. Therefore, in this hypothesized scenario, the central drive for accelerating theta and respiration would be the same. However, the changes in theta activity would precede changes in respiration rate, and thus look causal to them under directionality analyses, due to a presumed faster activation of theta-generating circuits, located a few synapses away, in comparison to the longer delays required for modulating respiration, which include modulation of the underlying networks, conduction delays to the diaphragmatic muscle, and subsequent thoracic volume changes.

**Figure 8.**
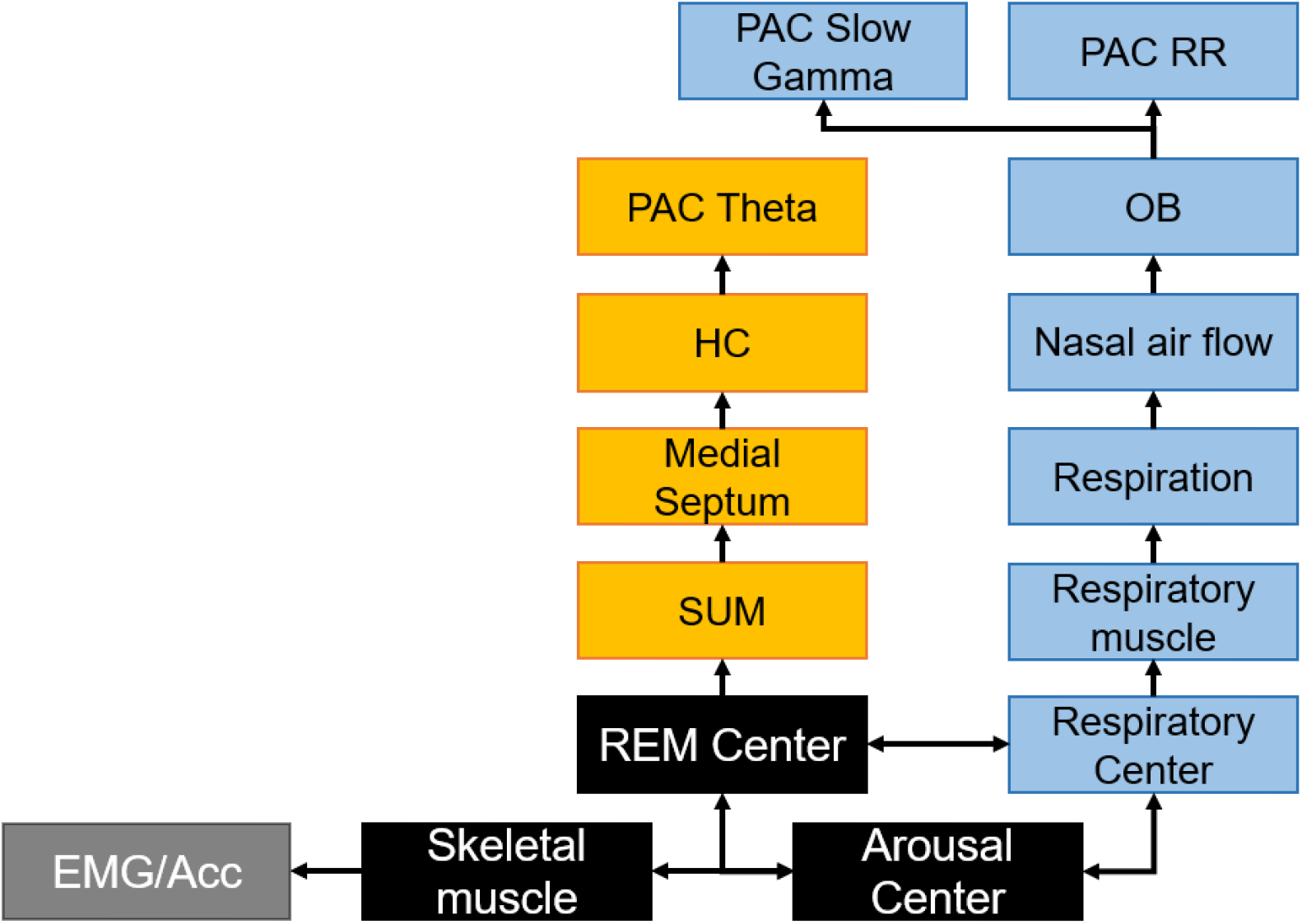
Schematics of hypothesized system interactions leading to the current results. See text for details. EMG: electromyography; Acc; accelerometry; SUM: supramammillary nucleus; HC: hippocampus; PAC: parietal cortex; RR: respiration-entrained rhythm.

Consistent with the model in Figure 8, there is experimental evidence for mutual connections among the brainstem microcircuits controlling REM sleep, arousal and respiration (Orem, 1980; Luppi et al., 2012; Boutin et al., 2017; Yackle et al., 2017; Benarroch, 2018; Del Negro et al., 2018). Interestingly, there are also connections between breathing centers and the generators of other orofacial rhythms such as whisking (Moore et al., 2013; Kleinfeld et al., 2014; McElvain et al., 2018), which may become active during REM sleep (Robinson et al., 1977; Tiriac et al., 2012). Therefore, the interplay among brainstem nuclei is likely to control higher arousal states within REM sleep that would be associated with faster theta and respiration frequencies. The scenario of changes in arousal is compatible with the accelerometer signal also being Granger-causal to respiration rate (Figure S2). Even though animals were asleep and therefore with low muscular activity, the accelerometer signal may have captured twitches of the skeletal musculature that happen during REM sleep (Figure 8; Robinson et al., 1977; Tiriac et al., 2012) and/or breathing-related body movements.

A possibility is that these activity states play a role in processing emotional information, one of the main functions attributed to REM sleep (van der Helm et al., 2011; Groch et al., 2013; Hutchison and Rathore, 2015; Tempesta et al., 2018), especially considering that experiencing emotions has been related to changes in breathing patterns (Grossman and Wientjes, 2001; Philippot et al., 2002; Zhang et al., 2017). But irrespective of the information content during these states, a recent study has shown that the respiration rate is the main modulator of cerebral oxygenation (Zhang et al., 2019), pointing to a close link between breathing and metabolic brain demands. Therefore, the higher oxygenation brought about by faster breathing indicates greater energy consumption, suggesting network states of heightened information processing.

Regarding potential behavioral or cognitive consequences of changes in breathing rate, it is worth noticing that differences in nasal flow rate influence olfactory processing (Buonviso et al., 2006; Courtiol et al., 2011a, 2011b; Esclassan et al., 2012). Based on recent studies, it is well feasible that breathing rate changes could also influence non-olfactory processes (Zelano et al., 2016; Varga and Heck, 2017; Arshamian et al., 2018; Herrero et al., 2018). Interestingly, using the same dataset analyzed here, we have recently shown that theta-gamma coupling, a hallmark network activity previously associated with cognitive processes (Tort et al., 2009; Canolty and Knight, 2010; Lisman and Jensen, 2013), does depend on the breathing rate through an inverted V-shaped relation (Hammer et al., 2020). This novel result suggests that indeed changes in breathing may influence non-olfactory information processing. As a technical remark, we note that the phenomenon of theta-gamma coupling could not be subjected to the same directionality analyses as investigated here for the individual oscillations. This is because assessing cross-frequency coupling requires using longer epoch lengths (i.e., sampling several theta cycles to average out noise; Tort et al., 2010; Scheffer-Teixeira and Tort, 2018) and cannot be performed in the same sub-second time windows used in the present study to assess the instantaneous frequency. In addition to influencing theta-gamma coupling, another recent study indicated that the breathing rate may also affect LFP phase-entrainment by respiration (Girin et al., 2020; see also Jessberger et al., 2016), suggesting a possible interplay between the rate and phase-driven effects.

Strikingly, while the instantaneous breathing rate followed changes in theta (Figure 3), the same directionality analyses revealed that respiration was causal to slow-gamma activity (Figures 6 and 7). This result suggests that neocortical slow-gamma is likely to depend on the olfactory bulb activity, which is majorly driven by sensory stimulation of the olfactory epithelium (Buonviso et al., 2006; Courtiol et al., 2011a, 2011b; Esclassan et al., 2012; Short et al., 2016). It is worth noticing that this drive is also provided by the mechanical stimulation through nasal air movement even in the absence of odors (Grosmaitre et al., 2007; Connelly et al., 2015). The olfactory bulb is well-known to express prominent gamma activity (Ravel et al., 2003; Martin et al., 2004; Beshel et al., 2007; Cenier et al., 2008, 2009; Kay et al., 2009), and, as a matter of fact, this region has been shown to produce more than one type of gamma with different sensitivities to respiration (Kay, 2003; Zhong et al., 2017; Zhuang et al., 2019). Interestingly, original accounts on gamma activity related this rhythm to olfaction (Adrian, 1950; Bressler and Freeman, 1980; Rojas-Líbano and Kay, 2008), even when recorded outside primary olfactory regions (Vanderwolf, 1992, 2001). While the present study measured gamma over the parietal neocortex, it would be interesting to study its relation to olfactory bulb gamma. Indeed, a recent report indicates that olfactory bulb gamma is causal to gamma in the medial prefrontal cortex (Karalis and Sirota, 2018). Moreover, Kay and Freeman (1998) reported a decrease in olfactory bulb low (but not high) gamma power during sniffing in an odor identification task, which could relate to the decrease in genuine parietal slow-gamma power with breathing rate observed here (Figures 4 and 5).

As opposed to slow gamma, genuine (1/f-corrected) fast-gamma power was not as causally related to the respiration rate (Figures 6 and 7). It is worth highlighting that the parietal recordings exhibited a clear excess power in the fast-gamma range (a “power bump”, see Figure 4A). Moreover, similar to previous characterizations of neocortical activity during REM sleep (Scheffzük et al., 2011, 2013; Brankačk et al., 2012), theta-fast gamma coupling was prominent in the present dataset (Hammer et al., 2020). In other words, even though the power levels at the fast frequencies of the LFP spectrum may reflect unspecific, aperiodic activity including EMG contamination, the current data did exhibit a true oscillatory activity in the fast-gamma range, that is, a genuine fast LFP rhythm. But despite the unequivocal presence, fast-gamma oscillations were not related to respiration, suggesting they could underlie internal computations not affected by respiratory reafference. Nevertheless, even though the individual oscillatory features were not affected, we have recently shown that the higher-order phenomenon of cross-frequency coupling between theta and fast-gamma does depend on breathing rate (Hammer et al., 2020), which underscores the complexity of respiration-brain interactions.

Previous studies in awake animals have shown that LFP activity in the theta and gamma ranges depend on running speed. The relationship between theta and speed is in fact an old one; since the first recordings, both theta power and theta frequency have been consistently shown to positively correlate with locomotion speed (McFarland et al., 1975; Sławińska and Kasicki, 1998; Hinman et al., 2011). More recently, speed-dependencies have also been reported for the gamma sub-bands (Chen et al., 2011; Ahmed and Mehta, 2012; Kemere et al., 2013; Zheng et al., 2015; Lopes-Dos-Santos et al., 2018). For instance, Chen et al. (2011), working with mice, reported a gradual amplitude increase with speed for both slow and fast gamma oscillations recorded in CA1. The same group later demonstrated changes in the instantaneous gamma frequency with speed in the rat CA1 (Ahmed and Mehta, 2012). Interestingly, this study also reported that slow gamma power negatively correlates with speed, a result subsequently corroborated by independent labs (Kemere et al., 2013; Zheng et al., 2015; but see Lopes-Dos-Santos et al., 2018). While we recorded from behaviorally immobile mice during REM sleep, the speed-dependencies of LFP oscillations described in these studies are reminiscent of the theta and gamma relations with the breathing rate observed here, especially considering that faster locomotion speeds are associated with faster breathing (Zhang et al., 2019). Future research that simultaneously records respiration and locomotion activity in suitable spatial arenas should help to disentangle the contribution of each factor to modulating LFP activity. In particular, from the present results we already know that changes in LFP activity with breathing may occur without locomotion changes. Likewise, it would be interesting to characterize the influence of speed after controlling for changes in breathing patterns.

Some technical aspects should be observed, though, such as the way power values are estimated, which may substantially vary among studies (i.e., absolute vs. relative power, correction or not for broadband power shifts, use of independent components or empirical, non-linear decompositions, etc; see Donoghue et al., 2020 for a recent discussion on this subject). As illustrated here, the precise method for estimating gamma power may critically influence the results (c.f. Figures 7 and S1). Another aspect is the recorded region (neocortex vs. hippocampus). While theta is considered a global rhythm, gamma represents more local activity and may thus exhibit different dynamics among regions. In fact, Zheng et al. (2015) showed differences in the speed-dependency of gamma activity between the entorhinal cortex and the hippocampus. To further complicate matters, Lopes-dos-Santos et al. (2018) called attention to potential differences in speed-modulations of hippocampal gamma activity between rats and mice. Finally, another technical aspect to be kept in mind is the very definition of gamma activity. For instance, the pattern which we call fast-gamma in the present work is particularly prominent in the neocortex during REM sleep (Scheffzük et al., 2011, 2013; Brankačk et al., 2012), and, despite the similar nomenclature, it is likely not to correspond to the ‘fast-gamma’ activity observed in the hippocampus of awake animals (Belluscio et al., 2012; Schomburg et al., 2014).

In summary, the present results show that the breathing frequency causally relates to theta and gamma network oscillations and that the relationship between respiration and neocortical brain activity goes beyond phase-entrainment relations. These findings motivate the investigation of breathing rate-relations in other brain regions, in different behavioral states and cognitive loads, and at different spatial scales, including the cellular level. Moreover, given the causal link between breathing and gamma shown here, it will be interesting to characterize further how specific sub-bands of gamma oscillations in different brain regions relate to olfactory bulb gamma and respiration.

## Materials and Methods

### Ethics statement

This study was approved by the Governmental Supervisory Panel on Animal Experiments of Baden-Württemberg (35-9185.81 G84/13 and G-115/14). All experiments were carried out in line with the guidelines of the European Science Foundation (2001) and the US National Institute of Health Guide for the Care and Use of Laboratory Animals (1996).

### Animal care

We used 22 C57/Bl6 mice (9 females and 13 males; age: 13-19 weeks; weight: 22-37 g). The animals were housed in groups of 4 with access to food and water *ad libitum*. After surgery for electrode implantation, animals were kept individually until completion of the experiments. Mice were housed on a 12-hour light-dark-cycle (lights off at 8:00 a.m., except 2 animals on the opposite circadian phase). Data from these same animals were used in a recent study of ours (Hammer et al., 2020).

### Surgery

Mice were implanted with epidural surface electrodes. Buprenorphine (0.1 mg/kg) was administered for pain control before and after surgery (every 8 h as necessary). Surgery was performed under isoflurane anesthesia (4% for induction, 1.5% during surgery; for further details, see Jessberger et al., 2016; Zhang et al., 2016). After skull exposure, 0.5 to 1 mm diameter holes were drilled above the parietal cortex according to stereotactic coordinates (2 mm posterior bregma, 1.5 mm lateral to the midline). Reference and ground electrodes were screwed into the skull above the cerebellum. Electrodes consisted of stainless steel watch screws. During surgery, body temperature was monitored and maintained at 37-38°C. Animals were allowed 7 days of recovery before experiments.

### Electrophysiological recordings and behavioral staging

Mice were placed in a whole-body plethysmograph (EMKA Technologies, SAS, France), which was customized to allow simultaneous recordings of local field potentials (LFPs). It consisted of a transparent cylinder (78 mm inner diameter, 165 mm height) connected to a reference chamber (Figure 1A). Individual recording sessions lasted from 2 to 6 hours (mean: 3 h). Monopolar electrophysiological signals were filtered (1-500 Hz), amplified and digitized at 2.5 kHz (RHA2000, Intan Technologies, Los Angeles, USA). A three-dimensional accelerometer was custom-mounted on the amplifier board located on the head of the mice to allow movement detection. The three signals of the accelerometer were fed to three channels of the amplifier using the same band-pass filter as for the LFPs (see above), therefore removing the gravity-induced sustained potentials of the accelerometers.

REM-sleep was manually staged by a senior researcher in the field (J.B.) and identified as follows: 1) minimal accelerometer activity and 2) continuous theta rhythm in the parietal cortex following a sleep stage with slow-waves typical for non-REM sleep (Brankačk et al., 2010).

### Data analysis

We used built-in and custom-written routines in MatLab^©^ (The Mathworks Inc., Natick, MA). For each animal, all identified REM sleep epochs were first detrended and then concatenated before analysis.

### Power and coherence

Respiration and LFP power spectra were computed using the pwelch.m Matlab function from the Signal Processing Toolbox. Phase coherence spectra between raw respiration and LFP signals were computed using the mscohere.m function (Signal Processing Toolbox). For both functions, we used 4-s windows with 50% overlap and nfft=2^16^.

Time-frequency power spectrograms were computed employing the spectrogram.m function (Signal Processing Toolbox). For the analyses shown in Figures 2, 4 and 5, we used 500-ms windows with no overlap. The numerical frequency resolution was set to 0.05 Hz for slower frequencies (0.5 to 20 Hz) and 0.2 Hz for faster frequencies (25 to 200 Hz). The power spectra shown in Figure 2C were obtained from the spectrogram in Figure 2A (mean over two 500-ms windows). For each 500-ms window, the corresponding peak frequency for theta or respiration was taken as the frequency with maximal power. For estimating slow and fast gamma peak frequencies, the power spectrum was first smoothed (15-Hz moving average) and a 1/f fit was subsequently subtracted (see Figure 4A; Brankačk et al., 2012; Scheffzük et al., 2013). The absolute slow and fast gamma power analyzed in Figure S1 were taken as the mean power in the corresponding frequency range; the “1/f-corrected” (or “1/f-detrended”) slow and fast gamma power analyzed in Figures 6 and 7 were obtained as the mean power at the same frequency ranges but computed after subtracting the 1/f fit. For the directionality analyses shown in Figures 3, 6 and 7 (see subsection below), the time series of instantaneous frequency and power were estimated from spectrograms computed using 500-ms windows but with 90% overlap (i.e., 50 ms step or 20 Hz sampling).

### Directionality

For estimating directionality relations, the respiration and LFP raw signals (Figure 1) or the time-series of instantaneous frequency and power (Figures 3, 6 and 7) were first z-scored. Directionality was estimated by means of cross-correlograms (CCGs) and Granger causality. CCGs were obtained using the xcorr.m function with a maximal lag of 1 second, using the ‘coeff’ option to normalize CCG values between −1 and 1 (equivalent to the correlation coefficient). Granger causality was computed through the Multivariate Granger Causality Matlab Toolbox (MVGC; Barnett and Seth, 2014). In Figure 1B, raw LFP and respiration signals were down-sampled to 250 Hz prior to computing Granger causality. The Variational Autoregressive (VAR) model order was fixed as 20 for all analyses. After computing VAR model parameters (tsdata_to_var.m) and associated auto-covariances (var_to_autocov), the pairwise conditional spectral Granger-causality was obtained through the MVGC toolbox function autocov_to_spwcgc.m. The overall, “time domain” Granger-causality was obtained from integration (average) of the Granger spectrum through the function smvgc_to_mvgc.m.

### Statistical analysis

Data are expressed as mean ± SEM unless otherwise stated. The significance of the difference between the mean Granger causality in each direction was assessed using paired t-tests; one sample t-tests were used to infer the significance of CCG lags against 0 ms. Repeated measures ANOVA was used to assess for significant relations between LFP frequency or power and respiration frequency. Statistical significance was set at alpha = 0.05.

## Acknowledgments

This work was supported by the Deutsche Forschungsgemeinschaft (SFB 1134/A01; Dr 326/10-1), Bundesministerium für Bildung und Forschung (German-Brazil Cooperation grant No. 01DN12098), the Brazilian National Council for Scientific and Technological Development (CNPq), the Brazilian Coordination for the Improvement of Higher Education Personal (CAPES), and the Alexander von Humboldt Foundation. The funders had no role in study design, data collection and analysis, decision to publish, or preparation of the manuscript.

## Supplementary Material

Content: 3 Supplementary Figures + Legends

**Figure S1.**
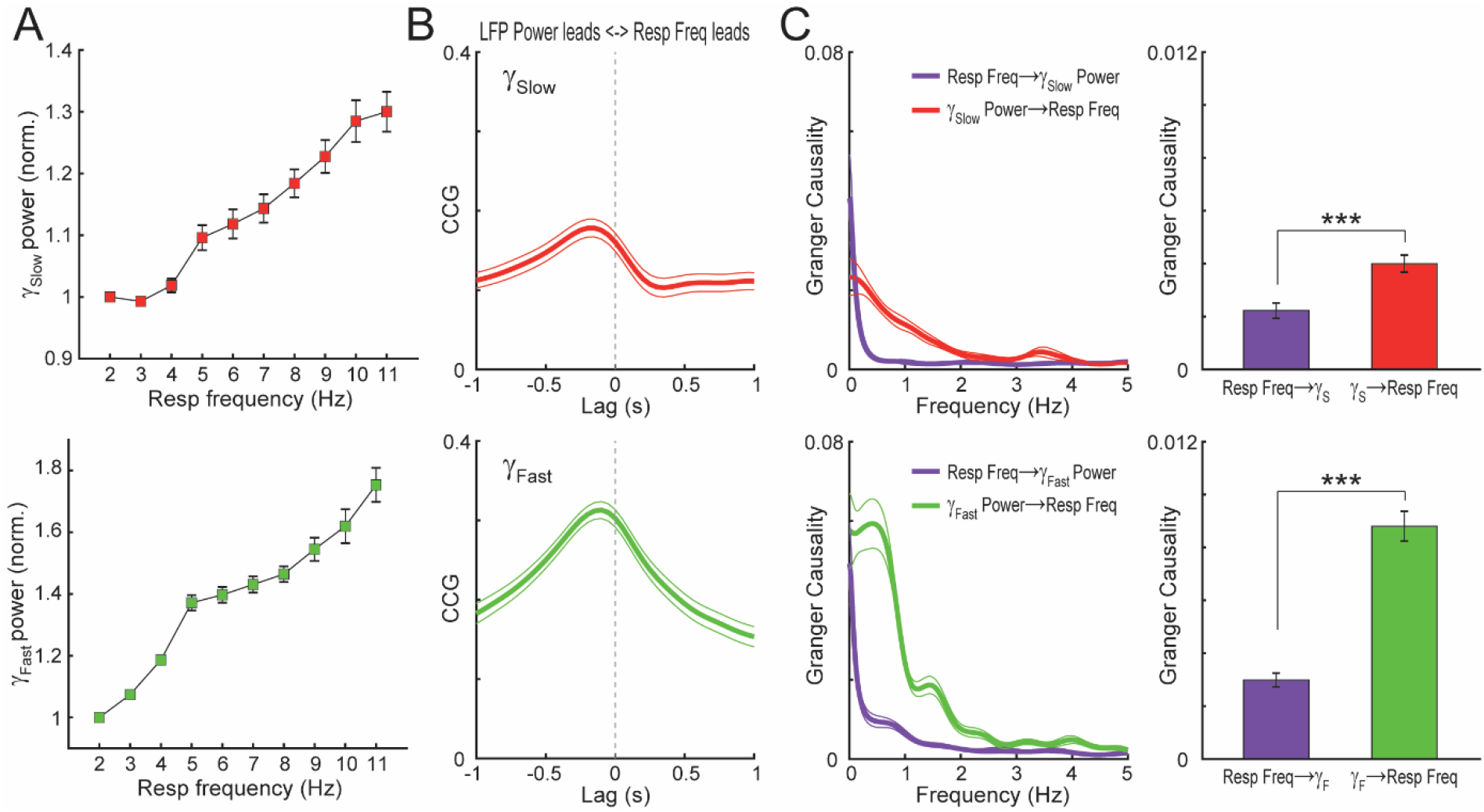
Absolute gamma power increases with breathing rate and is causal to it. (A) Group data of absolute gamma power as a function of respiration frequency (mean ± SEM; n=22 mice) for slow (top) and fast (bottom) gamma. Power values were normalized by the first respiration frequency bin. (B) Mean CCG (± SEM) between the absolute power of slow (top) and fast (bottom) gamma and respiration rate during REM sleep. (C) Frequency- (right) and time-domain (left) Granger-causality between absolute gamma power and respiration. Absolute power changes Granger-causes respiration frequency. ***p<0.001.

**Figure S2.**
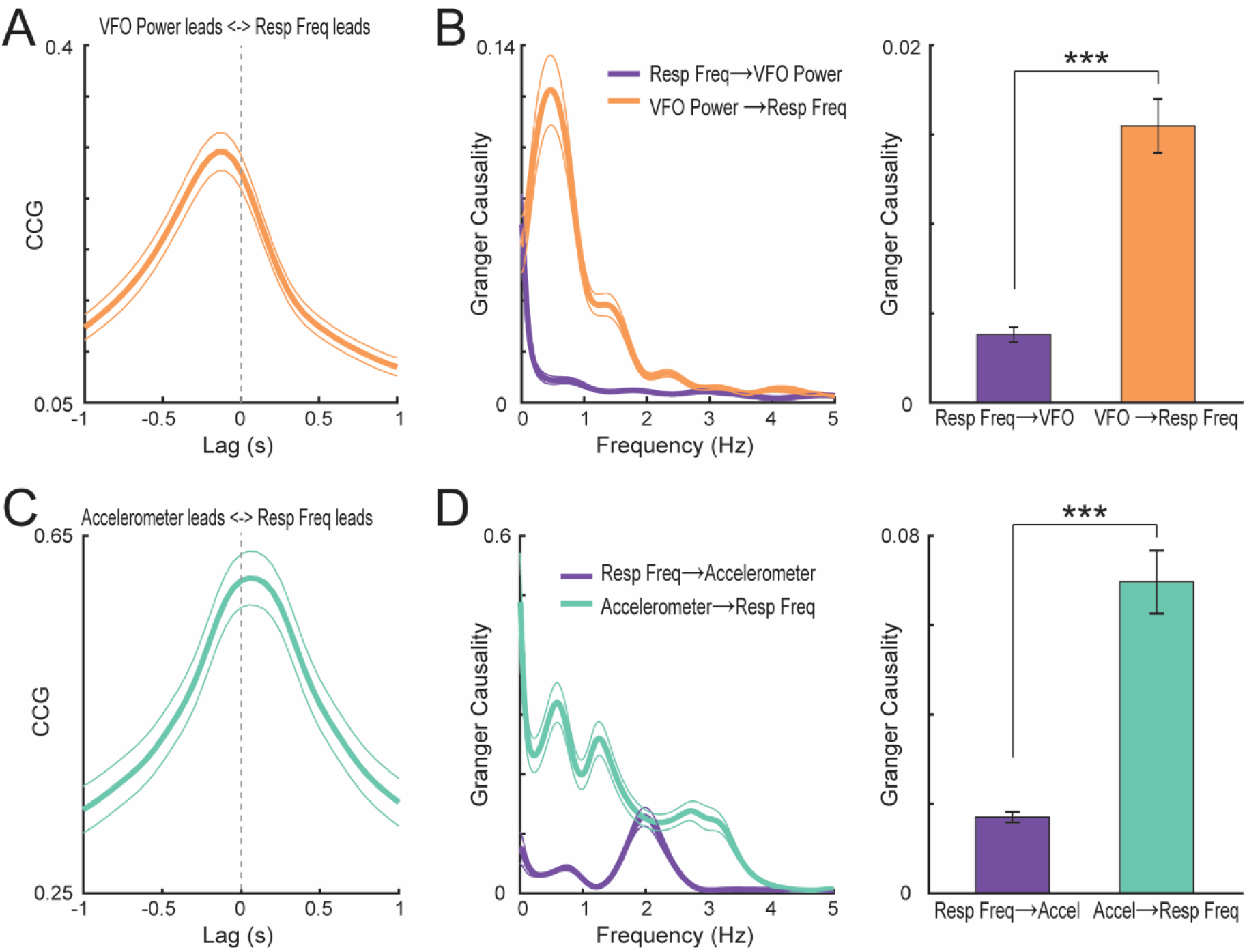
Both very-fast LFP power and accelerometer signals are causal to respiration changes. (A) Mean CCG (± SEM) between the absolute power of very fast oscillations (VFO, 180-230 Hz) and respiration rate during REM sleep (n=22 mice). (B) Frequency- (right) and time-domain (left) Granger-causality between VFO power and respiration. (C,D) Same as above, but for the root mean square of 3D accelerometer signals recorded at the headstage. Both VFO power and head acceleration during REM sleep highly Granger-cause respiration frequency. ***p<0.001.

**Figure S3.**
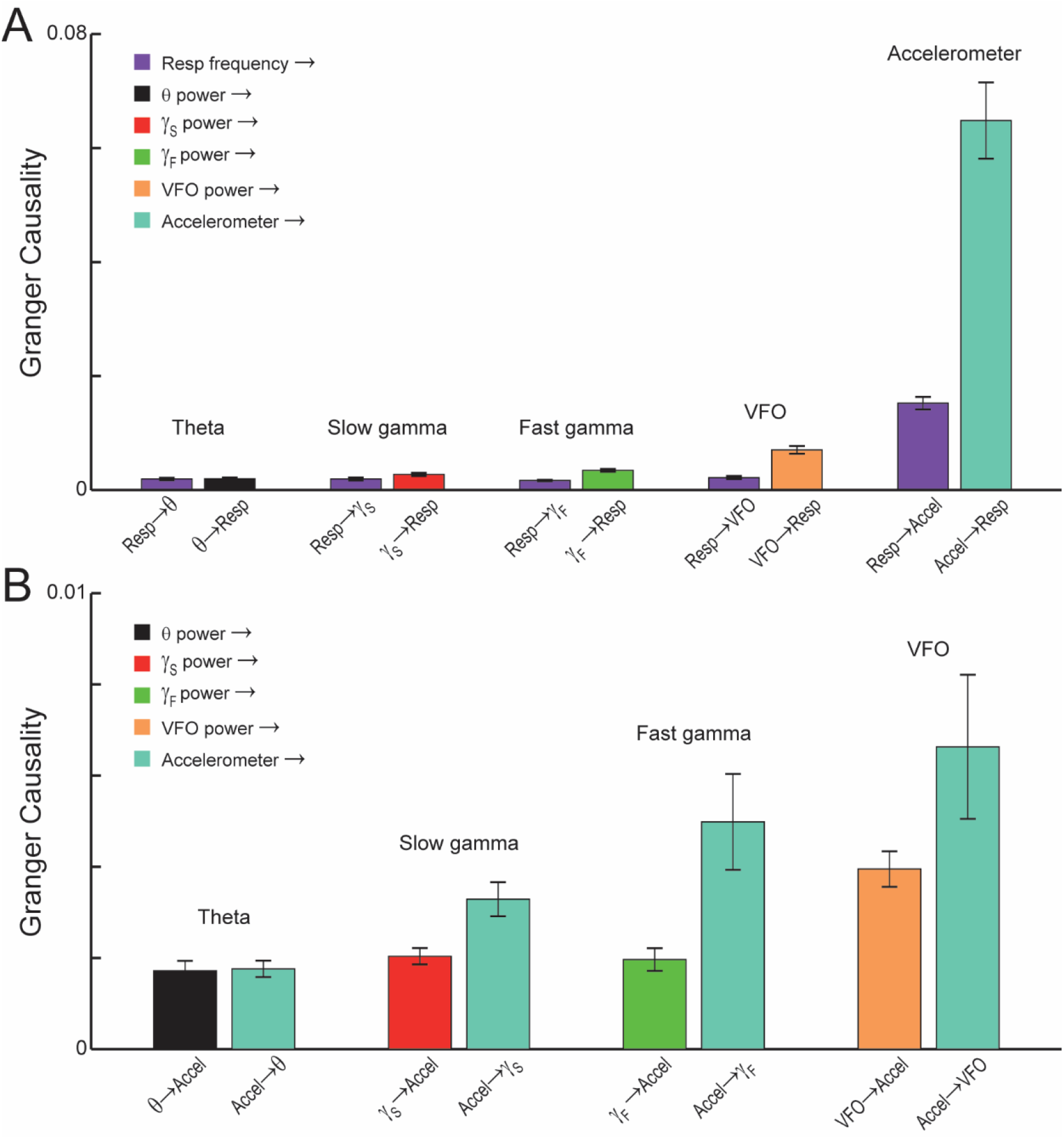
Accelerometer signals Granger-cause absolute power at fast LFP frequencies. (A) Granger-causality values for pair-wise combinations of respiration frequency and either LFP band power or the RMS of accelerometer signals. (B) Pair-wise Granger causality relations between head acceleration and LFP power.

